# Practical application of indicators for genetic diversity in CBD post-2020 Global Biodiversity Framework implementation

**DOI:** 10.1101/2022.02.18.481087

**Authors:** Henrik Thurfjell, Linda Laikre, Robert Ekblom, Sean Hoban, Per Sjögren-Gulve

## Abstract

Genetic diversity is a key aspect of biological variation for the adaptability and survival of populations of species, which must be monitored to assure maintenance. We used data from the Swedish Red-List 2020 and a recent government report to apply three indicators for genetic diversity proposed for the post-2020 Global Biodiversity Framework of the Convention on Biological Diversity (CBD). We made more detailed indicator assessments for mammals and herptiles.

For indicator 1, the proportion of populations with effective population size N_e_>500, 33% of 22557 investigated species had a population estimate. For herptiles and mammals, 70% and 49%, respectively likely had N_e_>500.

For indicator 2, the proportion of populations or range remaining within species, 20% of all species evaluated for the Red-List have data. Meanwhile, 32% of the herptile and 84% of the mammal populations are maintaining their populations/range.

For indicator 3, the number of species/populations in which genetic diversity is monitored using DNA-based methods, there are studies on 3% of species, and 0.3% are monitored. In contrast 68% of mammals and 29% of herptiles are studied using DNA, and 8% of mammals and 24% of herptiles are genetically monitored.

We conclude that the Red List provide data suitable for evaluating the genetic indicators, but the data quality can be improved. There is a synergy in estimating the genetic indicators in parallel with the Red-Listing process. We propose that indicator values could be included in national Red-Listing as a new category - “genetically threatened”, based on the genetic indicators.

## Introduction

The Convention on Biological Diversity (CBD; www.cbd.int) has from its ratification in 1993 identified genetic diversity - biological variation within species - as one of three pillars of biodiversity. However, implementing the CBD with respect to genetic diversity has long lagged behind the other biodiversity pillars of species and ecosystems, particularly for wild species (Laikre 2010; Laikre et al. 2010). The initial draft from the CBD for a post-2020 biodiversity framework also largely neglected genetic diversity (CBD/WG2020/2/3 January 2020). In a letter to Science, Laikre et al. (2020) argues for all species to be genetically monitored in order to maintain their genetic diversity and evolutionary potential. Hoban et al. (2020) develops this framework and defines three genetic indicators, suitable for CBD reporting: 1) the number of populations with effective population size (Ne) above versus below 500, 2) the proportion of populations (or geographic range) maintained within species, 3) the number of species and populations in which genetic diversity is monitored using DNA-based methods.[2] Further elaboration of these proposed indicators have been provided in subsequent work, including the suggestion to use the census population size, Nc>5000 as a proxy for Ne>500 when the Ne/Nc ratio is not known for the focal species; in other words to assume an Ne/Nc ratio of 0.1 (Hoban et al. 2021a; Hoban et al. 2021b; Laikre et al. 2021). Currently, the Ne-indicator is proposed as a Headline Indicator in the post-2020 CBD Global Biodiversity Framework, while the proportion of populations maintained-indicator is proposed as a Component Indicator and the DNA-based indicator is suggested as a Complementary Indicator (CBD/WG2020/3/3/Add.1; CBD/WG2020/2/INF/2;CBD/SBSTTA/24/3Add.1). Conservation genetics researchers such as the Coalition for Conservation Genetics are, however, arguing for making the proportions of populations maintained indicator also a Headline indicator and the DNA-based indicator a Component indicator (Kershaw et al. In press) (Kershaw et al., in press).

Several attempts are now being made to apply these indicators. In South Africa, Mexico, and Costa Rica primarily indicators 1 and 2 are being tested in pilot work (Melanie Munoz, Guido Alonso Saborio Rodriguez, Jessica da Silva, Alicia Mastretta-Yanes, pers.comm.). In Switzerland and Sweden, indicator 3 is being elaborated (Fischer and Litsosis 2022; Laikre et al. 2008b). Through a systematic review, the EU COST action G-BiKE is also developing a standardized summary for indicator 3 (Michael W. Bruford, pers. comm.).

Here, we report on our attempt to assess if national Red-List data in Sweden include sufficient information for applying the indicators 1 and 2 (the Ne-indicator and the proportion of populations maintained-indicator, Figure 1), using already collected data on 22571 species, subspecies and populations (henceforth species unless otherwise specified) gathered for the Swedish Red-List classification (Ahrné, Bjelke et al. 2020). Both indicator 1 and 2 can be calculated and reported in the absence of genetic data (Hoban et al. 2020; Hoban et al. 2021a; Laikre et al. 2021). Hoban et al (2020) and Laikre et al (2020) suggested that numerous data sources could have information for reporting on these indicators including species’ recovery plans or the Red List. The Red List is a globally recognized, standard, and rigorous inventory of conservation status of plant and animal species, using quantitative criteria to evaluate the extinction risk of species. We also assessed indicator 3 based on data from already published work on genetic monitoring in Sweden (Laikre et al. 2008a; Posledovich et al. 2021). The data from the Red Listing was chosen as it is the most complete dataset we had access to where a large amount of species has been evaluated thoroughly and consistently. Red List assessments often contain an estimate of the global (or in this case national) census size of a species, and sometimes also contain census size of individual populations or subspecies within species, and either number of, or trend in populations or geographic range. However it is not currently known how many Red List assessments contain this information and could be used for reporting on genetic diversity indicators. In the Swedish red listing process, known values are often filled out, while unknown values are left blank. However, for some groups of species values may also be left blank even though they could be estimated, as focus is usually on species that are more obviously rare or threatened.

**Figure 1.**
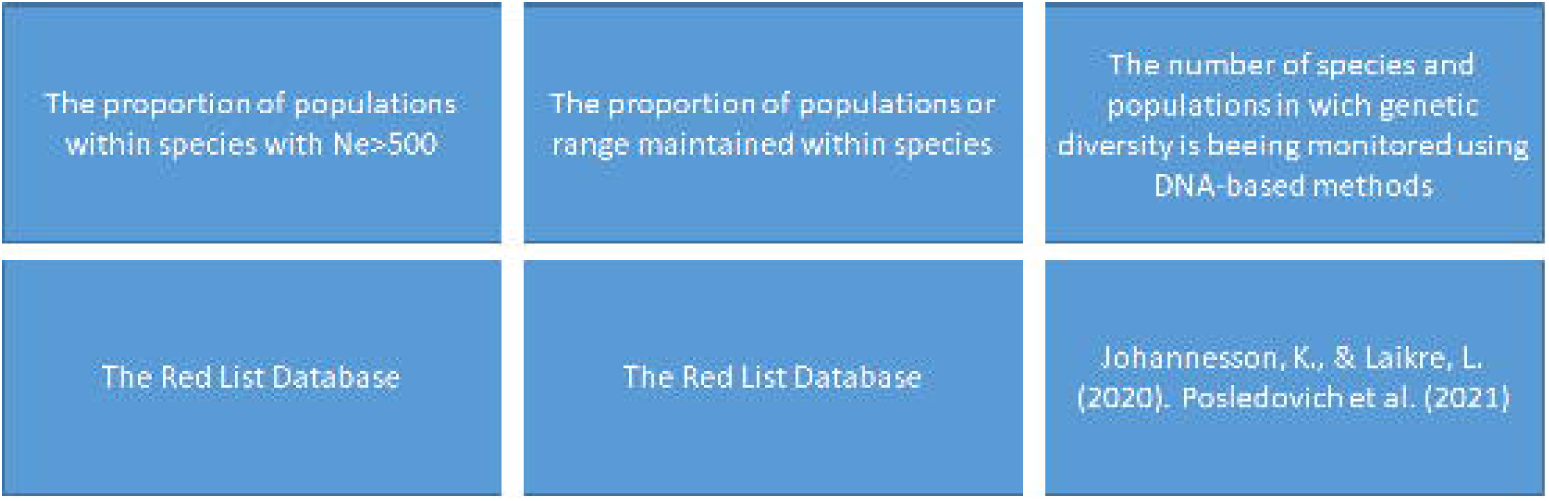
Genetic indicators and the source of information used for evaluation.

We assessed what proportion of the 22557 species that data on population(s) within species is available and how the data availability differs among species groups.

In an in-depth analysis, we selected two species groups from the Swedish Red List for which more detailed information is available, and calculated indicators 1, 2 and 3 for these species. For the selected groups, reproduction is sexual, there were relatively high-quality information, and they were (mostly) land bound terrestrial. These groups were mammals, amphibians and reptiles; amphibians and reptiles were combined into “herptiles” in the analysis (Ahlen et al. 2020).

## Materials and methods

We used Swedish Red-Listing data from a total of 22557 species that have been possible to classify [i.e., classified to either of categories, Least Concern (LC), Near Threatened (NT), Vulnerable (VU), Endangered (EN), Critically endangered (CR) or Regionally Extinct (RE)], to find out if and for how many species sufficient data is available for assessing proposed CBD indicators for genetic diversity (Hoban et al. 2020). Species that were Data Deficient (DD), Not evaluated (NE) or Not Applicable (NA), were excluded. Note that DD species are rare, and often it is not even certain they are reproducing consistently within Sweden.

In the Red List, only species that were present within the country in the year 1800 are assessed. Species that have immigrated on their own to Sweden are included if they can be regarded as established, which is classified to have occurred after 10 consecutive years of reproduction. Further, in a few cases known distinct subpopulations within species are Red-Listed separately (IUCN 2012). In this Swedish example, such listing applies to the porpoise (*Phocoena phocoena*, L.) harbour seal (*Phoca vitulina*, L.), red deer (*Cervus elaphus*, L.) and European grass snake (*Natrix natrix*, L.), that have distinct subpopulations which are genetically separated from the other populations (Ahrné et al. 2020) and these populations are assessed separately.

Generally, the Red Listing in Sweden is conducted by committees of experts on specific organism groups, each headed by a chairman employed at the Swedish Species Information Centre (https://www.artdatabanken.se/en/). The process follows an overarching standard but varies depending on the organism group, and how much information is available. For more well-known groups such as vertebrates, most fields that can be estimated are filled, while for many invertebrate groups, plants and fungi, a short-list of likely candidates for Red-Listing is made, and the focus is to estimate relevant values for those. The species not making the short-list are either common but unmonitored, or there is extremely limited knowledge. If there is no knowledge at all, they can be classified as data deficient, DD. The Swedish Red List has been updated every fifth year since 2000.

The following information that could be used for calculating the suggested genetic indicators is gathered per assessed species in the Red-Listing process (IUCN 2012): distribution trends and size of the total population, area of occupancy (AOO, number of 2*2km grid cells occupied by the species) and extent of occurrence (EOO, the minimum convex polygon or α-hull of species occurrences), degree of population fragmentation, size of the largest sub-population and if there is a negative trend in the number of localities or a negative trend in the number of sub-populations. If the species total population size within Sweden is assumed to be below 20,000 individuals, there is generally more effort focused on obtaining better estimates of population size than if the total population size is assumed to be above 20,000 individuals. The quality of the estimates varies greatly from best guesses of experts to very accurate monitoring data, but is generally best among well-studied groups such as vertebrates.

To calculate indicator 1, and its data availability, we classified all species with a value for total national population estimate, size of largest subpopulation or a classification for severely fragmented or Red Listed as RE, as having data. If any of these population estimates (national species census estimate or the size for the largest sub-population) were <5000, the species was classified as severely fragmented, or if it was red listed as RE, we classified it as having an Nc<5000 individuals.

In the more detailed analysis of mammals and herptiles, we analyzed data at the species level (e.g. we merged subspecies or subpopulations of the same species). The best population estimate was used to assess whether or not the species’ total population size in Sweden was larger than 5000 individuals. We assigned all species with a total population <5000 or Red Listed as RE and extinct in the last 100 years as 0% reaching the target Ne>500, as no subpopulation can be larger than the total population. One species that went extinct in Sweden more than 100 years ago was excluded from analysis (the wild reindeer). For species with known distinct (sub) population structure, the exact proportion of subpopulations with Nc>5000 was obtained. We then made some assumptions about remaining species based on their level of fragmentation and dispersal ability. For species where we don’t know the population structure, but we know the species are common and highly mobile, for example red fox (*Vulpes vulpes*), we assume that ≈100% of the populations of that species reach Ne>500 through gene flow. We then assessed the rest of the mammals and herptiles as follows. Rather common species that surely have some subpopulations on islands etc, such as the common frog (*Rana temporaria*) were given a value of “high”, which corresponds to 90% in the later summary calculations. For species classified as severely fragmented, we set the value either as “medium” (50%) or “low” (10%) depending on our knowledge of the species. A couple of species have been kept in game enclosures and escaped, been released, or translocated for hunting purposes, those populations are commonly<5000 individuals, but they hold little or no conservation value from a genetic standpoint, therefore they were assigned “anthropogenic” and excluded from the evaluation summary. Many of the Swedish red-deer populations are of introduced origin, but the nominate subspecies in Scania, the southernmost county of Sweden, is indigenous and its population is estimated to exceed 5000 individuals (Ahrné et al. 2020). We summarized the average proportions per species group using the above values.

To calculate indicator 2 - the proportion of populations within species maintained - all species that had any data filled out for any of the following criteria: decreasing extent of occurrence, decreasing area of occurrence, decrease in area or quality of habitat, or decrease in number of local areas or populations were considered to have data. If they were filled out as “yes” or red listed as RE they were classified as potentially losing populations.

For mammals and herptiles, we analyzed data on a species level, by merging separately assessed taxa of the same species. All criteria for trends in the red list are based on a three-generation time-span, therefore we also looked at a longer historical context of 100 years, and if the species had a larger distribution in the past, we classified it as not maintaining populations. Species red listed as RE that went extinct more than 100 years ago, were excluded from the analysis. Current population range compared to historical population range was estimated based on information in the Swedish species information database Artfakta (Artfakta 2022). Given the data in the Red-Listing database, it was hard to estimate “the proportion of populations maintained” within the species for most species (as indicator 2 is phrased), partly because data has only been recorded in a standardized manner since 2000, and here we have chosen a longer time horizon, so the response was mostly binary yes or no, for the summary calculated as 0% (no) 100% (yes). Many species were not assessed in the Red-List as losing or not losing populations, if they could not be assigned an exact value, they were estimated to ≈100% if they are good dispersers and likely not losing any populations, and “High” if they are common, but potentially some populations could have been lost. High corresponds to 90% in the summary.

To calculate indicator 3, we used already published work including reports from the Swedish Environmental Protection Agency to find which species had some genetic studies, and they should contain all studies up until 2020 (Johannsesson 2020; Posledovich et al. 2021). We combined the lists of all published studies in the scientific literature with all grey literature, and compiled it on a species level to assess all species with any study on population genetics. All studies with at least two time points (which could be considered monitoring), and all species with ongoing monitoring of genetic diversity were noted. Species where part of the population were studied was included, while species where studies are planned but not started were excluded (Table 1).

**Table 1.** Organism groups as they are evaluated per committee in the Swedish Red-List.

| Group | No. of species | Species with at least one population genetic study | Species for which at least one study including temporally separate studies of genetic diversity. | Species with regular monitoring of genetic diversity |
| --- | --- | --- | --- | --- |
| Mammalia | 69 | 47 (68%) | 29 (42%) | 5 (7%) |
| Aves | 261 | 60 (23%) | 11 (4%) | 24 (9%) |
| Herptilia (Reptilia and amphibia) | 21 | 17 (81%) | 6 (6%) | 5 (24%) |
| Pisces | 132 | 50 (38%) | 18 (14%) | 9 (7%) |
| Lepidoptera | 2614 | 27 (1%) | 2 (0.1%) | 5 (0.2%) |
| Hemiptera | 1024 | 1(0.1%) | 0 (0%) | 0 (0%) |
| Orthoptera | 35 | 3 (9%) | 0 (0%) | 2 (6%) |
| Coleoptera | 4281 | 22 (1%) | 1 (0.02%) | 0 (0%) |
| Hymenoptera | 982 | 14 (1%) | 4 (0.4%) | 0 (0%) |
| Odonata (and Plecoptera, Ephemeroptera, Trichoptera, Psochoptera) | 384 | 27 (7%) | 3 (1%) | 4 (1%) |
| Diptera | 1849 | 11 (1%) | 1 (0.05%) | 0 (0%) |
| Crustacea | 152 | 18 (12%) | 1 (1%) | 0 (0%) |
| Arachnida | 756 | 6 (1%) | 0 (0%) | 0 (0%) |
| Myriapoda | 71 | 0 (0%) | 0 (0%) | 0 (0%) |
| Anthozoa | 41 | 1 (2%) | 0 (0%) | 0 (0%) |
| Annelida (and Tricladida) | 76 | 16 (21%) | 0 (0%) | 0 (0%) |
| Echinodermata | 57 | 0 (0%) | 0 (0%) | 0 (0%) |
| Tunicata | 31 | 0 (0%) | 0 (0%) | 0 (0%) |
| Brachiopoda | 3 | 0 (0%) | 0 (0%) | 0 (0%) |
| Mollusca | 446 | 30 (7%) | 2 (0.4%) | 0 (0%) |
| Fungi (Only Macrofungi) | 3477 | 34 (1%) | 0 (0%) | 0 (0%) |
| Lichenes (Composite organism) | 1375 | 8 (1%) | 0 (0%) | 0 (0%) |
| Algae | 412 | 13 (3%) | 0 (0%) | 1 (0.2%) |
| Tracheophyta | 2985 | 149 (5%) | 3 (0.1%) | 2 (0.07%) |
| Bryophyta (and, Antho-cerotophyta, Marchantiophyta) | 1023 | 22 (2%) | 1 (0.1%) | 0 (0%) |
| Summary | 22557 | 576 (3%) | 82 (0.4%) | 57 (0.3%) |

## Results

### All species, indicator 1

A total of 7336 species (33%) of the 22557 assessed in the national red list in Sweden had some type of data on population size or structure (fig1a). Of those 7336 species 60% potentially have populations with Nc>5000 for indicator 1. (Figure 2a).

**Figure 2.**
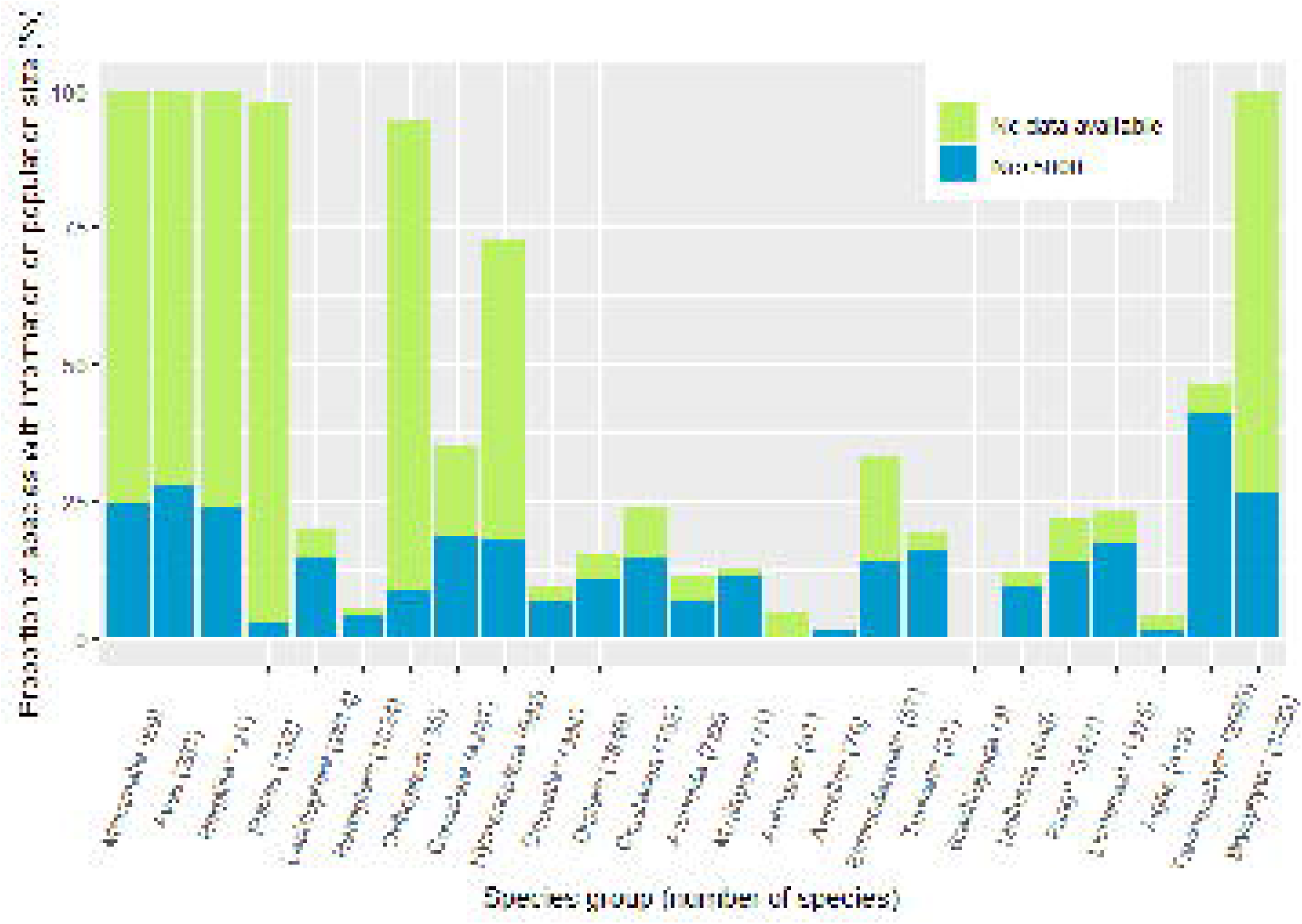
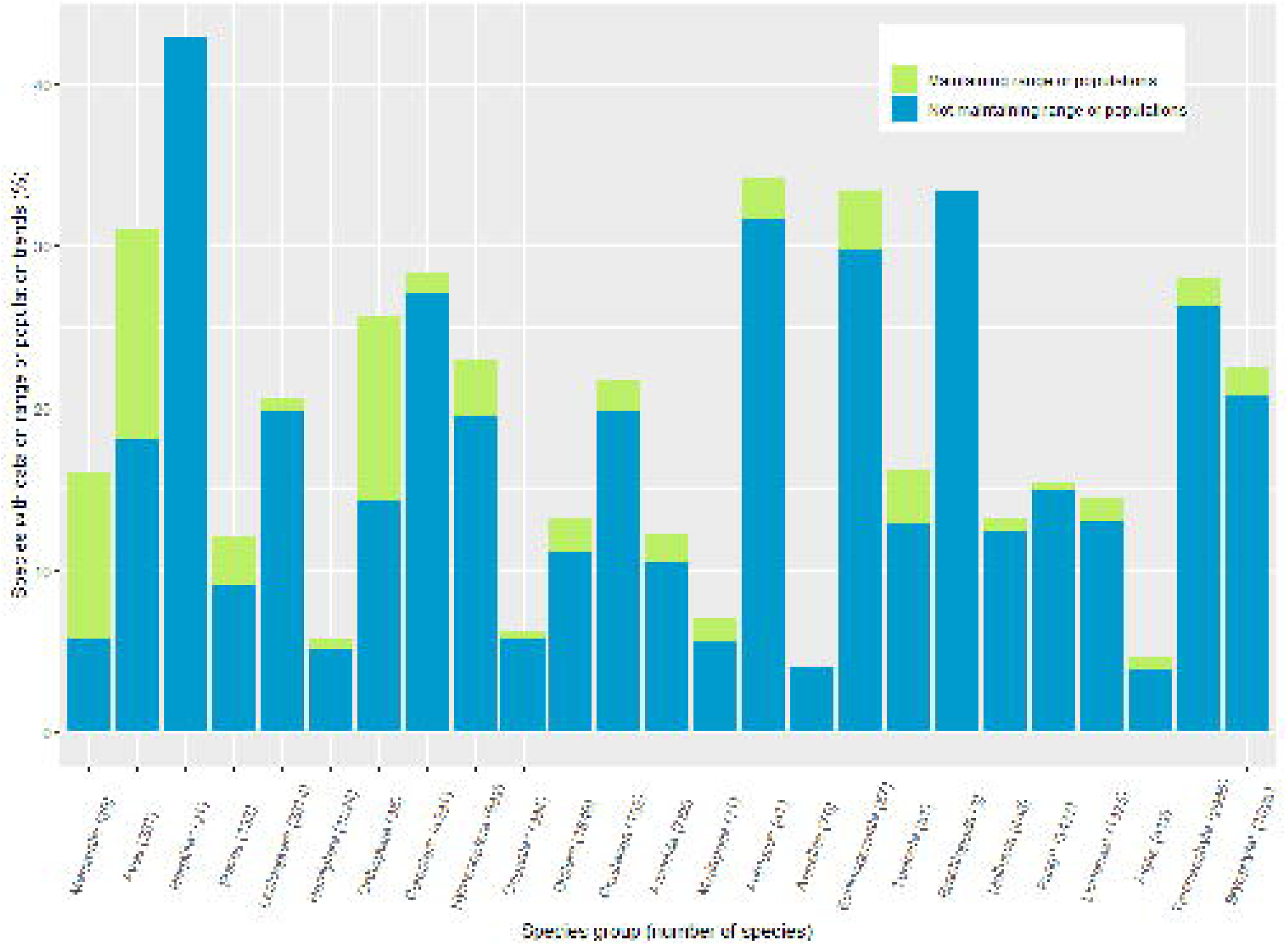
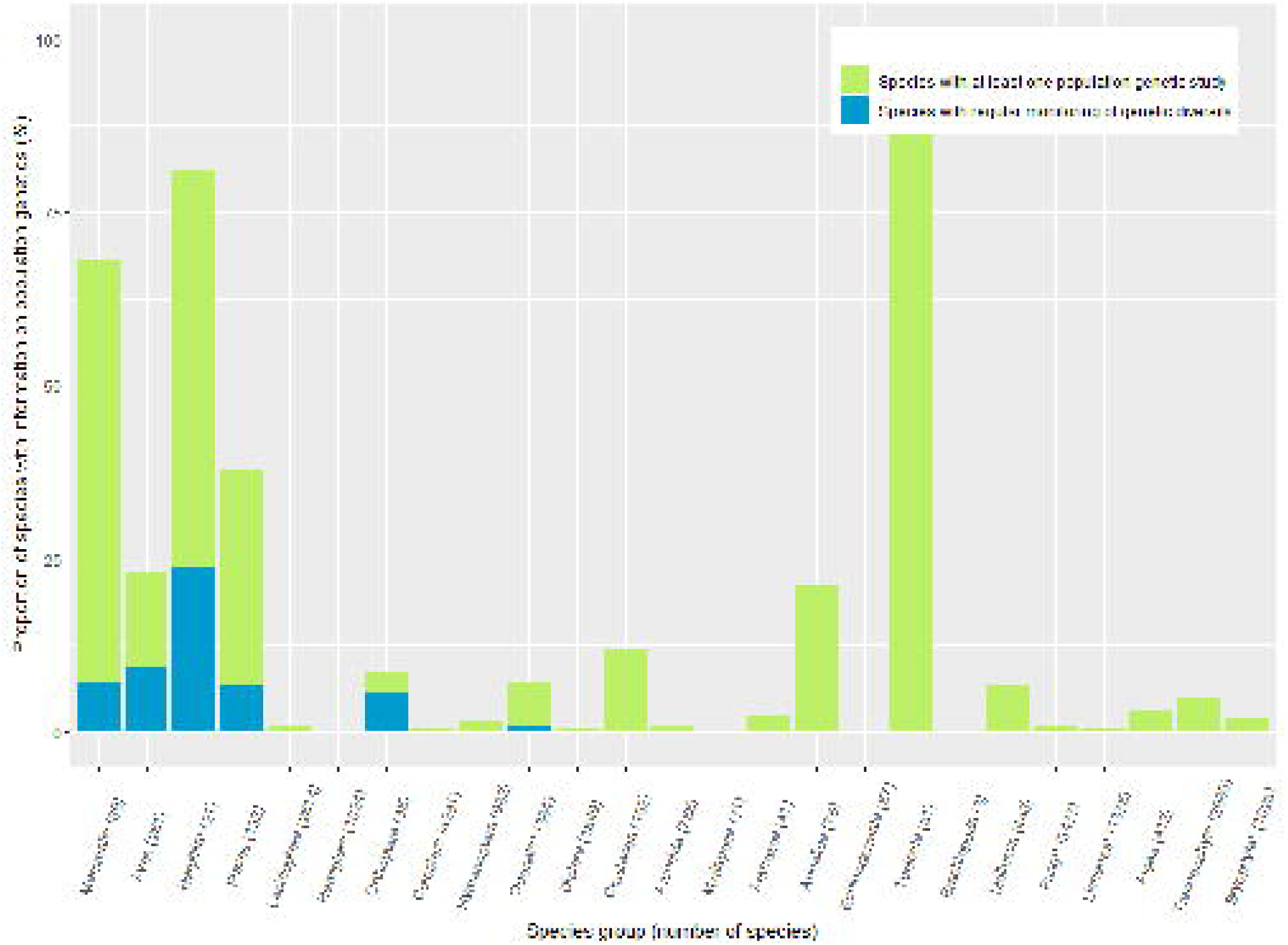
Available data for Genetic indicators per group. *See table 1 for definition of group. a. Indicator 1 b. Indicator 2 c. Indicator 3

Mammals and herptiles indicator 1 (Table 2); 49% of herptile and 70% of mammal populations in Sweden are estimated to be Nc>5000 individuals. All of them have a population estimate, or the population could easily be estimated to be >>5000 individuals by us. First, for 19 (30%) mammal species and 6 (35%) herptile species, the exact proportions of populations with Nc>5000 could be calculated. Second, in 13 mammal and 1 herptile species, the exact number of populations was known, the rest can be calculated due to a low total population. Third, for another 19 (30%) mammal species, very accurate proportions of populations with Nc>5000 could be estimated given the total population>5000 individuals and high mobility of species likely leading to a continuous national population. Fourth, for 23 (36%) mammals and 8 (47%) herptiles, the proportion of populations with Nc>5000 was estimated based on our knowledge of the groups.

**Table 2.** Species of mammals and herptiles assessed for the genetic indicators. Exact values comes from studies or deduction. ≈ was given to very good estimates. A written “Yes” or “No” comes from the red-list (for indicator 1 or 2), while “High”, “Medium” or “Low” were estimated. Summaries for indicator 1 and 2 were calculated as averages, where High was set to 90%, medium 50% and low 10%. For indicator 2, “Yes” (from the red list) was set as 100% and “No” 0%. For indicator 3 information on whether genetic diversity is monitored using DNA based techniques was obtained from Posledovich et al. (2021) their Appendix 3.^1^Known population structure (through research and/or deduction). ^2^Most red deer populations are of a mixed, anthropogenic origin, but there are two populations of the native subspecies (*C. elaphus elaphus)*, the values for those are in here. The anthropogenic populations of *C. elaphus* and *D dama* are excluded from analysis.

| Scientific name | Group | Indicator 1<br>% of pop<br>>5000 | Indicator 2<br>Maintains range or<br>populations | Indicator 3<br>Monitoring genetic diversity<br>with DNA-based techniques |
| --- | --- | --- | --- | --- |
| *Sorex isodon* | Mammals | 0% | No | No |
| *Sorex araneus* | Mammals | High | High | No |
| *Sorex minutus* | Mammals | High | High | No |
| *Sorex minutissimus* | Mammals | High | High | No |
| *Sorex caecutiens* | Mammals | High | High | No |
| *Neomys fodiens* | Mammals | High | High | No |
| *Canis lupus* | Mammals | 0%<sup>1</sup> | 100% | Yes |
| *Erinaceus europaeus* | Mammals | High | High | No |
| *Talpa europaea* | Mammals | High | High | No |
| *Lynx lynx* | Mammals | 0%<sup>1</sup> | 100% | No |
| *Ursus arctos* | Mammals | 0%<sup>1</sup> | 100% | Yes |
| *Vulpes lagopus* | Mammals | 0%<sup>1</sup> | 0% | Yes |
| *Vulpes vulpes* | Mammals | ≈100% | ≈100% | No |
| *Phoca vitulina* | Mammals | 50%<sup>1</sup> | 50% | No |
| *Halichoerus grypus* | Mammals | 50%<sup>1</sup> | 50% | No |
| *Pusa hispida* | Mammals | 0%<sup>1</sup> | 0% | No |
| *Lutra lutra* | Mammals | 0%<sup>1</sup> | 100% | No |
| *Meles meles* | Mammals | ≈100% | ≈100% | No |
| *Martes martes* | Mammals | ≈100% | ≈100% | No |
| *Mustela erminea* | Mammals | ≈100% | No | No |
| *Mustela nivalis* | Mammals | ≈100% | Yes | No |
| *Mustela putorius* | Mammals | ≈100% | Yes | No |
| *Gulo gulo* | Mammals | 0%<sup>1</sup> | 100% | Yes |
| *Pipistrellus nathusii* | Mammals | ≈100% | ≈100% | No |
| *Pipistrellus pipistrellus* | Mammals | 0% | High | No |
| *Pipistrellus pygmaeus* | Mammals | ≈100% | ≈100% | No |
| *Nyctalus noctula* | Mammals | ≈100% | ≈100% | No |
| *Nyctalus leisleri* | Mammals | 0% | High | No |
| *Eptesicus serotinus* | Mammals | ≈100% | ≈100% | No |
| *Eptesicus nilssonii* | Mammals | ≈100% | ≈100% | No |
| *Myotis nattereri* | Mammals | ≈100% | ≈100% | No |
| *Myotis dasycneme* | Mammals | 0% | High | No |
| Myotis bechsteinii | Mammals | 0% | High | No |
| Myotis mystacinus | Mammals | ≈100% | ≈100% | No |
| Myotis daubentonii | Mammals | ≈100% | ≈100% | No |
| Myotis myotis | Mammals | 0% | High | No |
| Myotis brandtii | Mammals | ≈100% | ≈100% | No |
| Myotis alcathoe | Mammals | 0% | High | No |
| Plecotus auritus | Mammals | ≈100% | ≈100% | No |
| Plecotus austriacus | Mammals | 0% | High | No |
| Vespertilio murinus | Mammals | ≈100% | ≈100% | No |
| Barbastella barbastellus | Mammals | ≈100% | ≈100% | No |
| Muscardinus avellanarius | Mammals | High | High | No |
| Castor fiber | Mammals | 100%<sup>1</sup> | 100% | No |
| Lepus timidus | Mammals | High | No | No |
| Alces alces | Mammals | 100%<sup>1</sup> | 100% | Yes |
| Capreolus capreolus | Mammals | ≈100% | ≈100% | No |
| Cervus elaphus | Mammals | Ex.<sup>1</sup>, 50%<sup>2</sup> | Ex. 100%<sup>2</sup> | No |
| Dama dama | Mammals | Ex.<sup>2</sup> | Ex.<sup>2</sup> | No |
| Phocoena phocoena | Mammals | 67%<sup>1</sup> | 67% | No |
| Sciurus vulgaris | Mammals | High | High | No |
| Sicista betulina | Mammals | High | High | No |
| Mus musculus | Mammals | High | High | No |
| Apodemus flavicollis | Mammals | High | High | No |
| Apodemus sylvaticus | Mammals | High | High | No |
| Rattus norvegicus | Mammals | High | High | No |
| Lemmus lemmus | Mammals | High | High | No |
| Myopus schisticolor | Mammals | High | High | No |
| Myodes rutilus | Mammals | High | High | No |
| Myodes glareolus | Mammals | High | High | No |
| Craseomys rufocanus | Mammals | High | No | No |
| Microtus agrestis | Mammals | High | High | No |
| Arvicola amphibius | Mammals | High | High | No |
| Alexandromys<br>oeconomus | Mammals | High | High | No |
| Summary Mammals |  | 70% | 84% | 8% |
| Triturus cristatus | Amphibians | medium | No | No |
| Lissotriton vulgaris | Amphibians | High | High | No |
| Bufo bufo | Amphibians | High | High | Yes |
| Bufo viridis | Amphibians | 0%<sup>1</sup> | No | Yes |
| Epidalea calamita | Amphibians | 0% | No | Yes |
| Rana dalmatina | Amphibians | High | No | No |
| Rana temporaria | Amphibians | High | High | Yes |
| Rana arvalis | Amphibians | High | High | Yes |
| Pelophylax lessonae | Amphibians | 0%<sup>1</sup> | No | No |
| Pelophylax esculentus | Amphibians | High | High | No |
| Pelobates fuscus | Amphibians | 0% | No | No |
| Hyla arborea | Amphibians | 0% | No | No |
| Bombina bombina | Amphibians | 0% | No | No |
| Lacerta agilis | Reptiles | Low | No | No |
| Coronella austriaca | Reptiles | medium | No | No |
| Natrix natrix | Reptiles | High | No | No |
| Vipera berus | Reptiles | High | High | No |
| Summary Herptiles |  | 49% | 32% | 29% |
| Total summary |  | 65% | 73% | 12% |

### All species, indicator 2

4470 (20%) of all species had data on if they were maintaining their populations, AOO or EOO or not, and 4142 (9%) had a potentially stable number of total populations or range for index 2 (Figure 2b).

### *Mammals and herptiles indicator 2* (Table 2)

32% of the herptile species and 84% of the mammals in Sweden have maintained their geographic subpopulations or distribution during the last 100 years.

### *All species, indicator 3* (Table 1)

There are population genetic studies on 576 species (2.5%) of all species in the Swedish Red List, 82 species (0.4%) have data from at least two time periods. There are genetic monitoring on 57 (0.3%). (Table 1, Figure 2c).

### Mammals and herptiles indicator 3 (Table 2)

There is at least one population genetic study on 68% of mammals and 81% of herptiles, 42% of mammals and 29% of herptiles have studies with at least 2 time points included that could be considered monitoring. There is ongoing genetic monitoring on 5 (8%) mammals and 5 (29%) herptiles (Table 2).

## Discussion

The inclusion of genetic diversity indicators in the CBD post 2020 framework is a major advance in a long-standing problem-the neglect of this vital aspect of biodiversity in policy (Hoban et al. 2020; Laikre et al. 2020). Still, genetic indicators require demonstration of their applicability, as they have not been used as much as species and ecosystem indicators, such as the Red List Index. The main difference between the genetic indicators and the Red List is a focus on different levels-while the Red List mainly focuses on the species level and avoiding species’ extinction, the genetic indicators are intended to look at a population level - genetic variation within and among populations and thus avoiding genetic extinction (Exposito-Alonso et al. 2021). The two perspectives are also derived from population ecology and evolutionary ecology, respectively, and are thus driven by different mechanisms, where species or populations may erode genetically before population effects can be seen (Kardos and Luikart 2021; Spielman et al. 2004). Even in a country with substantial environmental and species monitoring programs, it may be difficult to apply the genetic indicators to *all* organism groups, given current knowledge. On the other hand, the genetic indicators could be applied successfully to at least thousands of species overall per country, and high proportions of species in some taxonomic groups. From this test using Swedish Red-List data, the data availability is reasonably sufficient for mammals and herptiles (Table 1, Figure 2). Other groups that have reasonable data are fish, birds and some plants, although each presents it’s unique set of challenges. For freshwater fish, it is hard to assess connectivity between all different populations in each lake or waterway, for birds and marine fish a national assessment may be problematic as a large proportion of the populations may be transboundary, while plants have a large variation in the reproduction system.

There are two ways to view the results in Figure 2. First, we might conclude that we have data on a limited proportion of all species, and the effort to apply the Ne>500 across most or all species in a country would be extremely challenging. The other way to view the results is that good data exists when efforts have been made but no effort is currently invested into a population estimation for common species that are nowhere near to be Red-Listed. The high proportion of species with less than 5000 individuals in groups with a low proportion of species with population estimates suggests the latter approach has been applied to the lesser known groups (figure 2). However, it has been emphasized that population losses are ongoing in many species (Ceballos and Ehrlich 2002; Ceballos et al. 2017), and we should not neglect these population estimates for common species, even if they do not meet Red List criteria.

In the two more closely studied groups, it is possible to apply indicator 1 in better detail (proportion of species where Nc>5000) as most species in these groups have some population estimate, are known to be well over the threshold, and we have well-informed ideas on population structure for most species. No species with a population >5000 is listed as heavily fragmented or have a very limited area of occurrence or localities for these groups (Table S1). For other groups, we may have to adjust how these criteria are used, depending on population dynamics within the group. For instance, fragmentation does not mean the same thing for a clonal plant species, or species with a seed bank, as it does for an amphibian. Variation in how the Red Listing was approached may vary between the Red-Listing committees, therefore the best practice is likely that someone involved in the Red-Listing of the particular group is involved in the assessment of genetic indicators.

As we see in Figure 2, there is a huge variation among taxonomic groups both in the availability of data as well as in the proportion of species fulfilling Nc>5000. The reasons behind this vary greatly, as each group has its own individual challenges and opportunities. For the vertebrate groups (mammalia, aves, herptilia, pisces) efforts are usually made to evaluate population sizes regardless of whether the species will be Red Listed or not, and often data is available through monitoring programs and their importance for citizen science, hunting and fishing. However, the ecology of the species determines the main differences, as both species size, and landscape connectivity become important. Many herptiles have a range limited to the southernmost part of Sweden, limiting their total populations, while especially marine and brackwater fish for example, may have more continuous populations throughout their respective habitat. Such variation in data quality and quantity for red lists has been noted in other studies (Bland et al. 2012; Butchart and Bird 2010).

Data availability for Indicator 2 depends a lot on how each expert group approached the red-listing process. The more detailed study on mammals and herptiles shows it is definitely possible to apply the indicator, but a more in-depth evaluation of each species could render a better estimate instead of the current yes-no dichotomy (range declining or not) that can be derived from the Red-Listing data. The proxy of current distribution < historical distribution that we used for the more closely studied groups, requires an individual assessment that is not included in the Red-List, and hence time consuming for large groups. However, there is probably great synergy effects of making calculations for this indicator in parallel with the Red Listing process, as changes in habitat distribution and quality are often assessed for the species. For example, it contributed one species (*Hyla arborea*, L.) to the second indicator that would otherwise have been left out.

It is worth noting that from the Red Listing process, there is more abundant data for the second indicator, and it is likely more robust as population censuses are scarce. Interestingly there are several species, mainly Red-Listed under the B criterion (Limited and decreasing AOO or EOO), for example the Natterjack toad (*Epidalea calamita*, Laurenti) where loss of populations, or decrease in AOO or EOO have been estimated, based on knowledge of the species ecology, measured large scale habitat trends and to some extent citizen science data. The second indicator scales well, as the loss of individual populations can be recorded on most scales. It is also less dependent on the definition of individual/ census size, which can be tricky in many organism groups (especially some plants and insects). We also note there may be other data sources available other than the Red List which could be applied across large numbers of species, such as large species occurrence data, which has been used in some metrics of population loss, including a proposed indicator for plants (Khoury et al 2019; Powers and Jetz 2019)

For indicator 3, we used previous literature reviews and reports (Laikre et al. 2008b; Posledovich et al. 2021). The counting of genetic studies and genetic monitoring is not done within the Red-Listing process, but as species specific information is gathered for the Red-List, it would probably be a great synergy effect of doing these searches (e.g. searching for genetic studies) in parallel, e.g. during red list updates. Doing so can provide a benefit for the Red List by enabling a more accurate estimate of total population numbers or structures in some cases, only obtained from genetic information. Further, genetic monitoring data can be used to assess Ne (indicator 1) as well as more detailed information on the threat situation such as inbreeding assessments and direct estimates of rates of genetic change.

### Comparisons with the Red List

Compared to the Red List, the genetic indicators 1 and 2 indicate more species appear threatened under genetic criteria than under Red-List/ demographic criteria, given the higher threshold of 5000 instead of 2000 individuals for indicator one, and the longer time span 100 years instead of three generations for indicator 2. In particular this longer time scale will likely catch the depleted species that have plateaued at a low level with only a fraction of their former population. Depleted species are Red Listed as LC since their populations are stable for the last three generations if their population exceeds 2000 individuals, even though their population, for example, may only have 1% left. If those populations are investigated using a genetic indicator, the longer time span enables us to identify species that may be at risk, as genetic erosion may have happened (Exposito-Alonso et al 2021) and inbreeding in isolated populations may be a threat. However, there are a few more criteria in the Red-List and combinations of criteria that may lead to species being classified as NT or more threatened even if population size is >2000. In other words, genetic threats and Red List assessments are not aligned. The Red Listed species that did not meet genetic indicators in the more closely studied groups were highly mobile species such as bats, that may have a small fraction of a larger european population in Sweden, but also decreasing widespread common species. The genetic indicators 1 and 2 seem more sensitive to species with relatively low mobility such as herptiles, or species with a clear isolated population, such as marine mammals that are split into Baltic and Atlantic populations (Table S1).

### Further developments

The genetic indicators do not consider hybridization issues, where a native subspecies such as red deer (*Cervus elaphus elaphus*, L.), is being outbred with introduced animals of taxonomically mixed origin. For relatively well known species groups, these factors could be included in some way in an alternative genetic indicator, such as the genetic scorecard developed recently in Scotland (Hollingsworth et al. 2020).

One issue encountered when working with the genetic indicators proposed by Laikre et al (2020) and Hoban et al (2020) is the lack of publicly available, detailed guidelines so far for applying them in practice. This will need to be developed as assumptions may need to be made in order to apply them for now. For example in this study, the level of fragmentation was used to determine the proportion of populations might be below Ne 500, which we admit is a strong assumption. In addition, it was not possible to evaluate population structure for most species, hence the simplified dichotomy of ‘maintaining populations’ or ‘not maintaining populations’ so guidance on this aspect is needed. Without detailed guidance, there will likely be differences in results depending on who did the assessment, something that can reduce the credibility and repeatability of indicators over time. For the Red-List, definitions are well documented, and for the genetic indicators to take a more prominent role in conservation, definitions and guidelines needs to be developed.

We have a few suggestions. The first regards scale. A genetic indicator may be less well suited to assess at a country level if populations are transboundary; the best practice may be to do joint assessments on the appropriate scale in such case. However, for a population spanning several countries where a joint assessment is appropriate, there is also a joint responsibility for the conservation of the population. The second indicator seems less sensitive to spatial scales, as a loss of range or populations is always a loss of range or populations. However, a loss of range for a transboundary population may only be seen in some of the countries hosting the population. Second, guidance is needed on advice for defining populations, for different types of species, such as those with different mobility or dispersal ability. Third, a more complete database of Ne/Nc ratios could help apply a more species-tailored ratio than the assumed 0.1 ratio employed here.

### The next steps

When assessing species for the Red List, there are synergies if genetic assessment are done in parallel, as information is often gathered species by species and literature and experts for each species are consulted. Therefore, the easiest way to apply the genetic indicators to a larger proportion of species may be to do it at the same time as the Red-List is reassessed. The first step would be to instruct Red Listing committees to fill out all checkboxes that can be estimated, i.e. if “has a declining AOO” is not filled out as “yes”, it should be filled out as “no” if that can be estimated. The number of populations should always be filled out if possible. Another easy change is that for species where there is some knowledge on population size, there should be instructions to enter the population size as >20,000 (20,000 is the population size that can potentially affect the Red-Listing status in combination with other estimates) if that can be estimated. The next step would be to add info that is specific for the genetic indicators, such as the sizes of different populations, historical range estimates (maybe AOO and EOO for comparability) and information on genetic monitoring.

In addition, at this point, if there are species-specific estimates of Ne/Nc numbers, such ratios could be added instead of the approximation of Ne/Nc 0.1. Since such ratio estimates already are available for several species (Hoban et al. 2020; Hoban et al. 2021c), this would be an appropriate next step in the analysis of species in Sweden. Another easy modification would be to include population genetic experts in the evaluation teams.

A suggestion may be that genetic conservation indicators such as these could be implemented in the Red List, adding a new criterion, Genetically threatened. A similar suggestion to include a ‘genetic threat’ in the red list process has been made several times (Garner et al. 2020; Willoughby et al. 2015). We truly think that it is now feasible and necessary for the red list to do so, as the genetic threats to species are growing in impact, and as genetic technologies and knowledge of genetic processes are ever more accessible (Hoban et al. 2021b; Parli et al. 2021; Taft et al. 2020). Assessors could be asked to input the numbers of historic populations and current populations, and estimates of the census size of each population, in tables which would then be amenable to large scale analysis across many species.

Overall, we conclude that genetic indicators could be evaluated for a substantial fraction, but not a majority of species in a country, and that data quantity, and the indicator value, will differ for different taxonomic groups.

## Acknowledgements

This report was funded by the Swedish Environmental Protection Agency (SEPA), and they had a role in defining the questions, but not in the outcome.The work of LL was funded by the Swedish research council Formas (grant FR-2020/0008) and the Swedish Reseach Council (grant 2019-05503).

